# Real Time Estimation of Population Receptive Fields Using Gradient Descent

**DOI:** 10.1101/194621

**Authors:** Mario Senden

## Abstract

A real-time population receptive field mapping procedure based on gradient descent is proposed. Model-free receptive fields produced by the algorithm are evaluated in context of simulated data exhibiting different levels of temporally autocorrelated noise and spatial point spread. As with any model-free approach, the exact shape of receptive fields produced by the real-time algorithm depends on the stimulus. Nevertheless, estimated receptive fields show good correspondence with ground-truth receptive fields in terms of both position and size. Furthermore, fitting a parametric model to the previously obtained estimates approximates the exact shape of the true underlying receptive fields well.

## Introduction

Real-time estimation of population receptive fields (pRFs; ***Dumoulin and Wandell, 2008***) offers many practical advantages. It can, for instance, make population receptive fields immedediately available for subsequent real-time decoding experiments reconstructing perceived (or imagined) visual stimuli (***Senden et al., 2017***). It might also allow for online optimization of the mapping procedure, for example by guiding stimulus placement to obtain ideal coverage of the visual field. Here, a real-time population receptive field (pRF) mapping procedure is proposed. The procedure employs a gradient descent on weights connecting the visual field to the cortex. The resulting weights (reflecting population receptive field estimates) can be directly used for subsequent real-time experiments. Alternatively, they can provide the basis for fitting a parametric pRF model. Model-free (raw) pRFs are evaluated in light of simulated data stemming from a model of V1 topography in the context of different levels of noise and point spreads of the blood-oxygen-level-dependent (BOLD) signal. Evaluations occur in terms of the ability of the procedure to localize pRFs, approximate their shape, and the ability of these pRFs to explain variance in model BOLD activity in response to a separate test stimulus.

## Methods and Materials

### Stimulus Design

The stimulus is an important contributor to the accuracy with which population receptive fields can be estimated (***Senden et al., 2014***). In the present study, a novel multi-scale bar stimulus is used. The stimulus is split into three concentric regions. The inner region spans a diameter of 7° and is traversed in twelve discrete steps by a bar ∼0.58° in width. The intermediate region spans a diameter of 14° and is traversed in twelve steps by a bar ∼1.16° in width. The outer region spans a diameter of 21° and is traversed in twelve steps by a bar ∼1.75° in width. The upper row of ***Figure 1*** shows three representative stimulus configurations. As can be appreciated from the figure, bars in the three regions never overlap and can exhibit different orientations since they are presented in unrelated random sequences. Within each region, bars are presented in four different orientations with each orientation being presented six times. In total, there are 288 samples of stimulus presentation padded before and after by eight samples without stimulation leading to a total of 304 samples.

**Figure 1.**
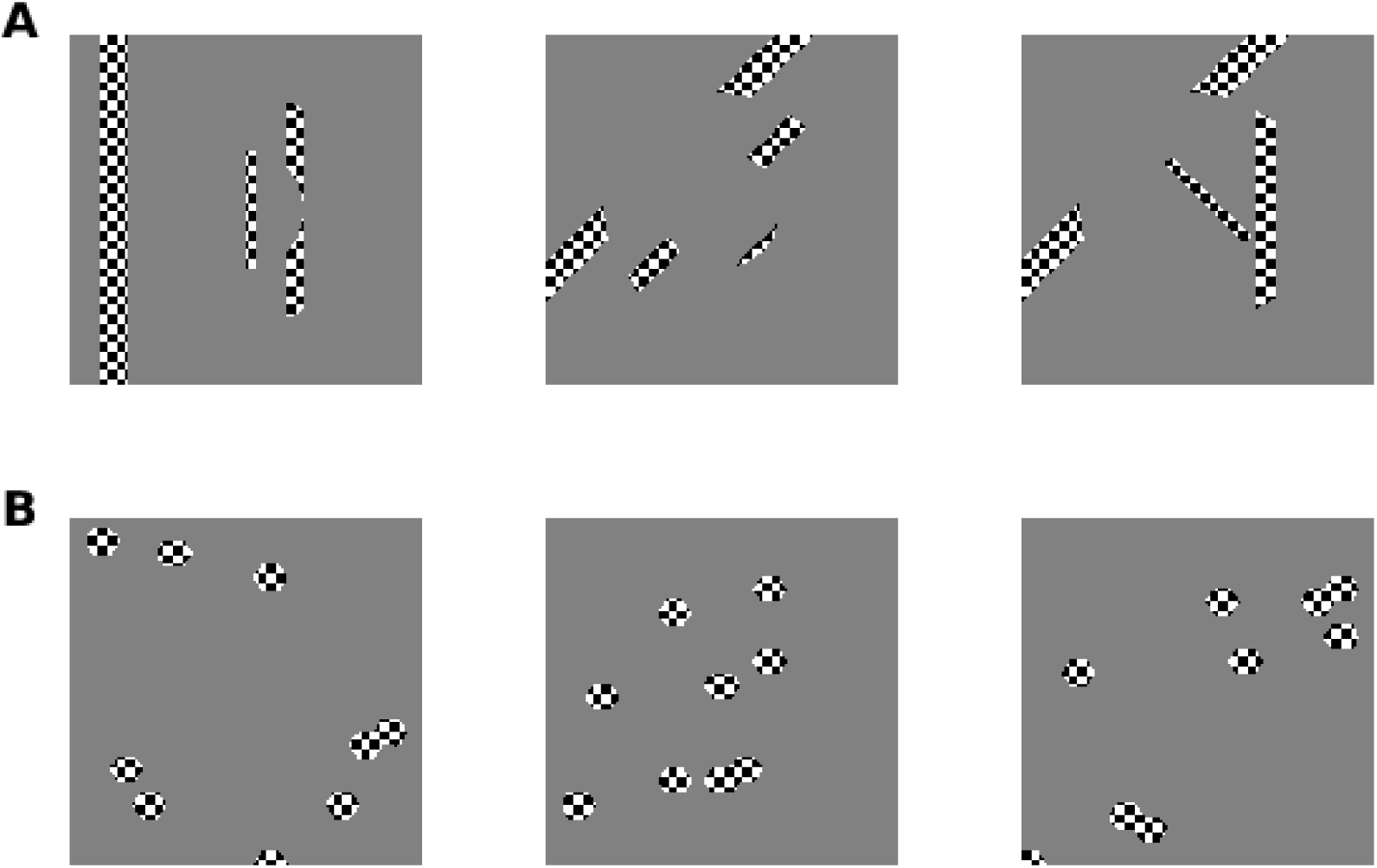
A) Three representative configurations of the multiscale bar stimulus. Bars are oriented independently at the three scales and presented with independent sequences. **B)** Three representative configurations of nine honeycomb blobs. Checkerboards are shown for illustrative purposes only. All simulations and mapping procedures are based on binarized effective stimuli.

In addition to the multiscale moving bar stimulus used for pRF mapping, a second stimulus is used to evaluate the predictive power of estimated receptive fields. This second stimulus consists of 288 random combinations of nine blobs 1.33° in diameter arranged in a honeycomb pattern. The lower row of ***Figure 1*** shows three representative stimulus configurations. Both stimuli have a resolution of 150×150 pixels.

## Model V1

### Cortical Surface and Receptive Field Properties

A V1-like cortical sheet extending 40 mm along and approximately 36 mm orthogonal to the horizontal meridian in both hemispheres consitutes the basis of the model. Since such a sheet is akin to a flattened cortical mesh, model units are referred to as vertices rather than voxels. Each vertex in the model is a 0.5 mm isotropic patch whose receptive field center is directly related to its position on the surface in accordance with a complex-logarithmic topographic mapping between cortical surface and visual field (***Balasubramanian et al., 2002***; ***Schwartz, 1980***) with parameter values (*a*=0.7°,*α* =0.9; ***Polimeni et al., 2005***). ***Figure 2*** shows the resulting eccentricity and polar angle maps across the cortical surface. The shape of model receptive fields is given by a 2-dimensional Gaussian

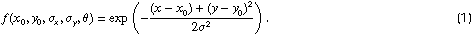

with (*x*_0_,*y*_0_) being the receptive field center and *σ* its size. Below an eccentricity of *E*=0.6° all model vertices have a receptive field size of *σ* = 0.125° whereas they exhibit a linear relationship with eccentricity (*σ* = 0.21*E*) beyond this cutoff (cf. ***Freeman and Simoncelli, 2011***).

**Figure 2.**
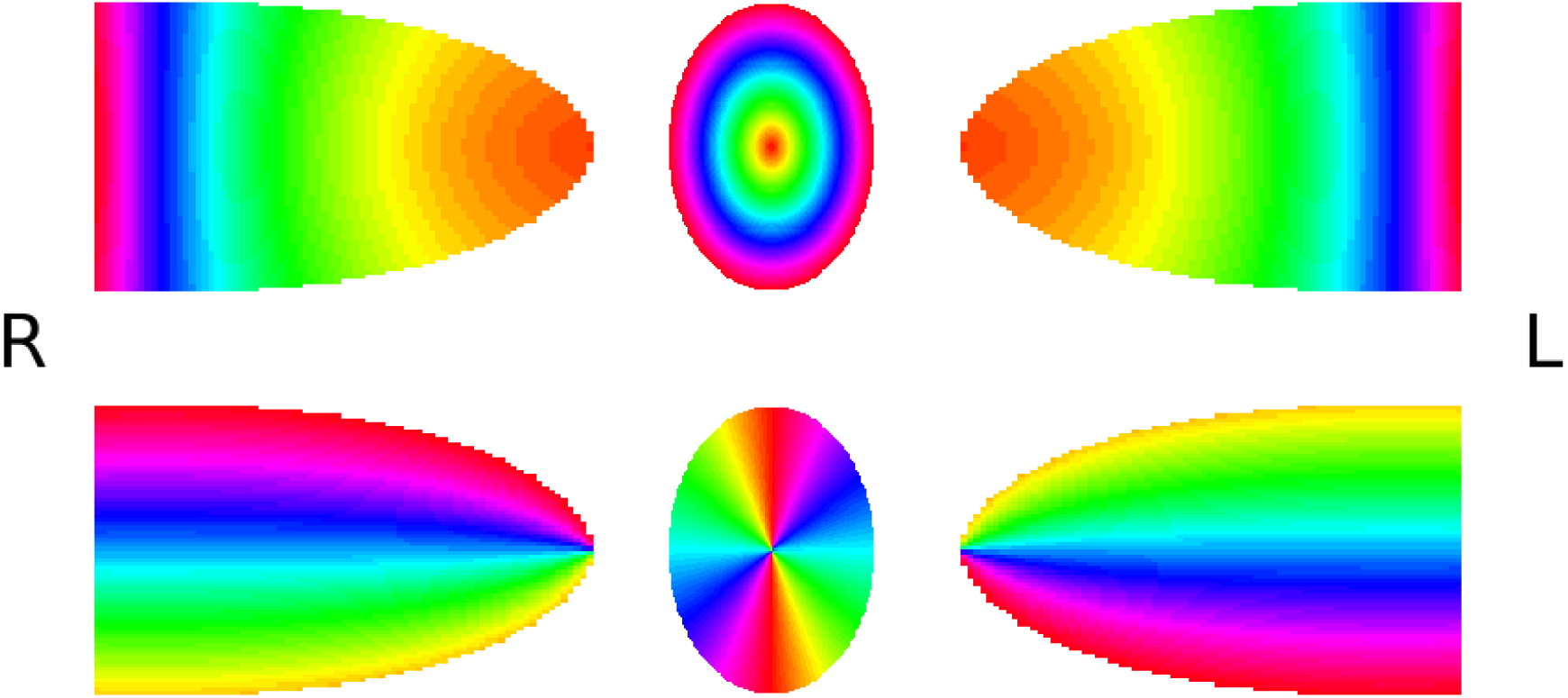
Eccentricity (top) and polar angle (bottom) maps representing the topographic organization of the V1 model. Following radiological convention, the left hemispheric surface map is shown on the right an vice versa. Centrally presented circles provide color codes for eccentricities and polar angles.

### Simulated BOLD Signal

A simulated BOLD signal (sampled at a rate of 0.5 Hz) for each vertex is obtained by first performing elementwise multiplication between the receptive field of a vertex and the effective stimulus presented per time point, summing the result, and subsequently convolving the thusly obtained timecourse with a hemodynamic response function. The hemodynamic response function of each vertex is given by a two-gamma function

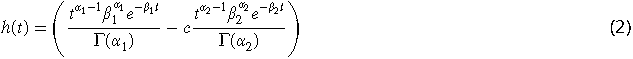

with parameters *α* and *β* controlling the shape and scale, respectively, and *c* controlling the ratio of response to undershoot. For each vertex, the five parameters are drawn from normal distributions around their canonical values. Specifically *α*_1_ ∼*𝒩*(6,0.5), *α*_2_ ∼*𝒩*(16,0.5), *β*_1_ ∼*𝒩*(1.1,0.1), *β*_2_ ∼*𝒩*(1.1,0.1), and *c*∼*𝒩*(0.167,0.1).

Two sources of distortion are added to the signal. First, a spatial smoothing kernel is applied to simulate the point-spread function of BOLD activity on the surface of striate cortex (***Shmuel et al., 2007***). Second, noise following a second-order autoregressive AR(2) model is added (*w*_*t*-1_ = 0.8,*w*_*t*-2_ = -0.2). The smoothing kernel is independently applied to the clean signal and the noise before the two are combined. The proposed mapping procedure is evaluated for a range of point spreads (2.25 mm to 4 mm in 8 steps) and SNRs (0.25 to 2 in 8 steps). Furthermore, performance results are evaluated in more detail (including fitting a parametric model) for a point spread of 2.5 mm full-width at half-maximum (FWHM; ***Shmuel et al., 2007***) and a signal-to-noise ratio 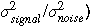 of 1.25.

## pRF Mapping Procedure

### Gradient Descent

The receptive field of each individual vertex *i* is considered to be a weight row vector **W**_*i*_ of length *p* connecting all pixels in the visual field to the vertex. The weight vectors of all vertices can then be combined into the *v*-by-*p* weight matrix **W**. At any given moment in time, the predicted BOLD activity (ŷ_t_) given this matrix and a column vector representing the effective effective stimulus **s**_**t**_ is then given by **y**_**t**_ = **Ws**_**t**_ and the discrepancy between the predicted and observed (z-normalized) BOLD activity (**y_t_**) can be measured by

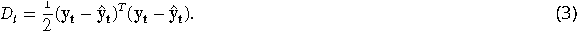

The gradient of the discrepancy with respect to the weight matrix is given by

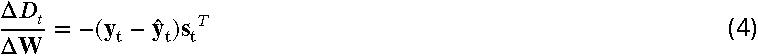

leading to the update of the weight matrix against this gradient according to

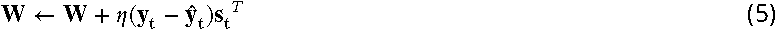

where η is a learning rate (1×10^-5^). The effective stimulus matrix **S** (of dimension *p*-by-*t*) is the result of convolving the timecourse of each pixel with the canonical two-gamma hemodynamic response function and z-normalizing the result. Weights are thus updated online with each time point constituting a new training example. Before training, initial weights are drawn from a normal distribution **W**∼*𝒩*(0,1×10^-5^).

### Real Time Signal Processing

In order for the described gradient descent approach to be used in a real-time setting, the effective stimulus needs to be convolved with a hemodynamic response in real-time. Furthermore, both the BOLD signal and the effective stimulus need to be z-normalized in real-time. Real-time convolution is achieved via a running convolution algorithm based on fast Fourier transformation (FFT) using an overlap-add scheme (***Wefers, 2015***). Specifically, as a preparatory step, the hemodynamic response function of length *N* is transformed using an FFT. Subsequently at each point in time *t*, the incoming signal of length 1 is zero-padded to length *N* and likewise transformed. The transformed signal and hemodynamic response are then multiplied elementwise and the result transformed back to the time domain by an inverse FFT. The final convolved signal is the result of adding up these overlapping partial convolution results (*x*_partial_):

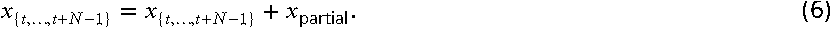

For real-time z-normalization, both mean and variance of the signal are estimated using online algorithms:

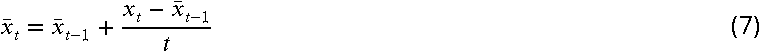

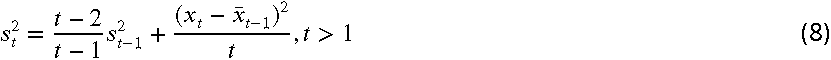

to subsequently obtain the z-score

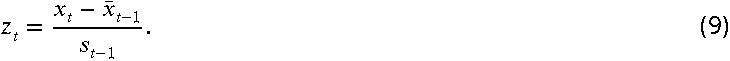

### Parametric Model Fitting

A parametric receptive field model can be fit to the estimated weight vector **W**_i_ of each vertex *i*. Here, in accordance with the ground-truth receptive fields, the parametric model is an isotropic Gaussian with parameter set *ψ* = (*x*_0_,*y*_0_,*σ*). Using the Levenberg-Marquardt algorithm (***Levenberg, 1944***; ***Marquardt, 1963***) implemented in MATLAB’s (2015a, The MathWorks, Natick, MA) optimization toolbox, the parameter set *ψ* for each vertex is found that solves the problem

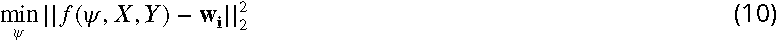

where *f* is the Gaussian function (cf. ***Equation 1***) and *X* and *Y* are x- and y-coordinates of each pixel in the visual field. The initial conditions *ψ*_0_ for each vertex are the x- and y-cordinate corresponding to the peak value in its weight vector and *σ*_0_ = 0.1. To improve the fitting procedure in light of potentially noisy raw pRF estimates, each weight vector **W**_*i*_ is squared and subsequently normalized with respect to its maximum value before applying the Levenberg-Marquardt algorithm.

## Results

### Mapping Performance

The proposed gradient descent approach to pRF mapping is evaluated for a V1 model with circular receptive fields. The performance of the algorithm is evaluated in terms of how well it localizes receptive fields as measured by the correlation between model and estimated eccentricity as well as polar angle maps, in terms of how well it reproduces receptive fields as measured by the correlation between model and estimated weights, and finally in terms of how well the BOLD activity predicted by the estimated weights for a novel stimulus protocol correlates with model BOLD activity observed for the same stimulus. ***Figure 3*** shows the performance of the algorithm as a function of SNR and point spread. For all considered measures SNR affects performance whereas point spread does not, at least within the considered range.

**Figure 3.**
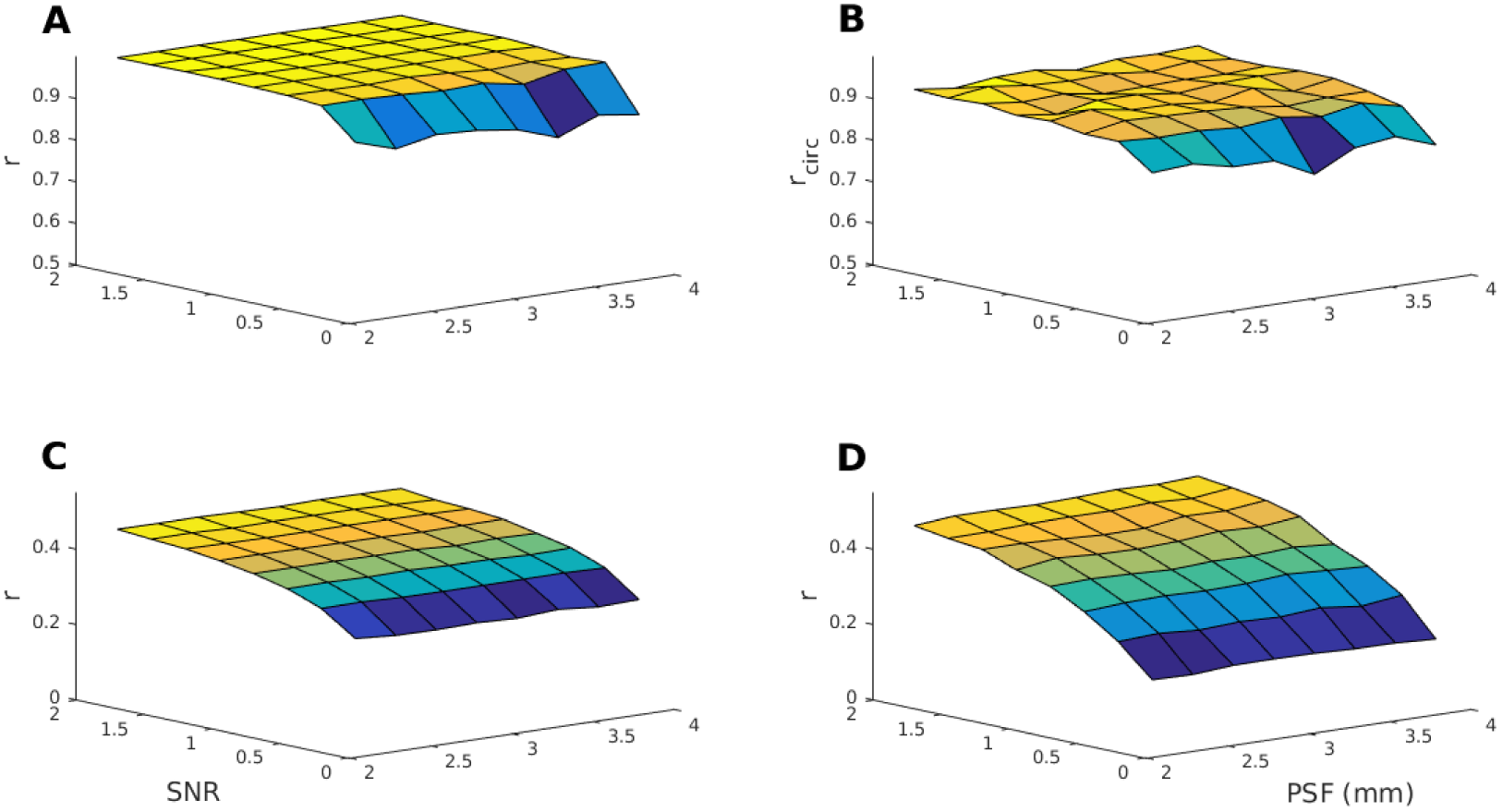
Pearson’s correlations between predicted and model pRFs and BOLD activity as a function of point spread and signal to noise ratio. **A)** Correlations between estimated and model eccentricity. Except for SNRs below 0.75, estimated and model eccentricity maps are virtually identical regardless of spatial point spread. **B)** Circular correlations between estimated and model polar angle. Except for SNRs below 0.75, estimated and model polar angle maps agree very well ∼ 0.9. Point spread has a negligible effect. **C)** Correlations between estimated and ground-truth receptive field shapes (weights). The estimated and model pRF shapes show good agreement. Agreement improves with higher SNR but is hardly affected by point spread. **D)** Correlations between estimated and model BOLD activity obtained for the test stimulus (see ***Figure 1***). As for the shapes, the estimated and model BOLD activity show good agreement. Agreement improves with higher SNR. This is expected since the model BOLD activity in response to the test stimulus become less noisy with increasing SNR. Point spread has again a negligible effect.

Next, a more in depth evaluation is presented for SNR = 1.25 and a point spread equal to 2.5 mm. ***Figure 4*** shows eccentricity and polar angle maps obtained from the gradient descent approach. The agreement with the underlying maps shown in ***Figure 2*** is visually striking. Indeed correlations between the model and estimated maps are *r*=0.98 and *r*_*circ*_ =0.91 for eccentricity and polar angle, respectively. ***Figure 5*** shows three representative ground-truth (panel **A**) and estimated (panel **B**) receptive fields at different eccentricities (and hence of different sizes). Correlations between model and estimated pRFs are *r*=0.15, *r*=0.48, and *r*=0.68, at the three eccentricities respectively. These results suggest that correspondence improves with receptive field size. ***Figure 6*** shows the correspondence between ground-truth and estimated receptive fields as a function of model pRF size. Panels (**A,C**) show the correlation between model and estimated receptive fields for raw estimates stemming directly from the gradient descent (**A**) and after fitting an isotropic Gaussian (**C**). Panels (**B,D**) show the correlation between BOLD activity in response to the test stimulus based on ground-truth and estimated pRFs, again for raw estimates (**B**) and after fitting an isotropic Gaussian (**D**). As can be appreciated from the figure, correlations for raw estimates increase with receptive field size. This is likely due to non-zero residual weights covering a larger proportion of the visual field for small as compared to large receptive fields. Consequently, the relationship between correlation and receptive field size is reduced after fitting an isotropic Gaussian to the raw estimates. At the same time, correlations are generally larger both for receptive fields and predicted BOLD activity. Indeed, correlations between groundtruth and estimated pRFs for the three previously described receptive fields increase to *r* = 0.59, *r* = 0.98, and *r* = 0.97, respectively, after fitting a parametric model. Finally, ground-truth and estimated receptive field sizes correlate well with *r*=0.95.

**Figure 4.**
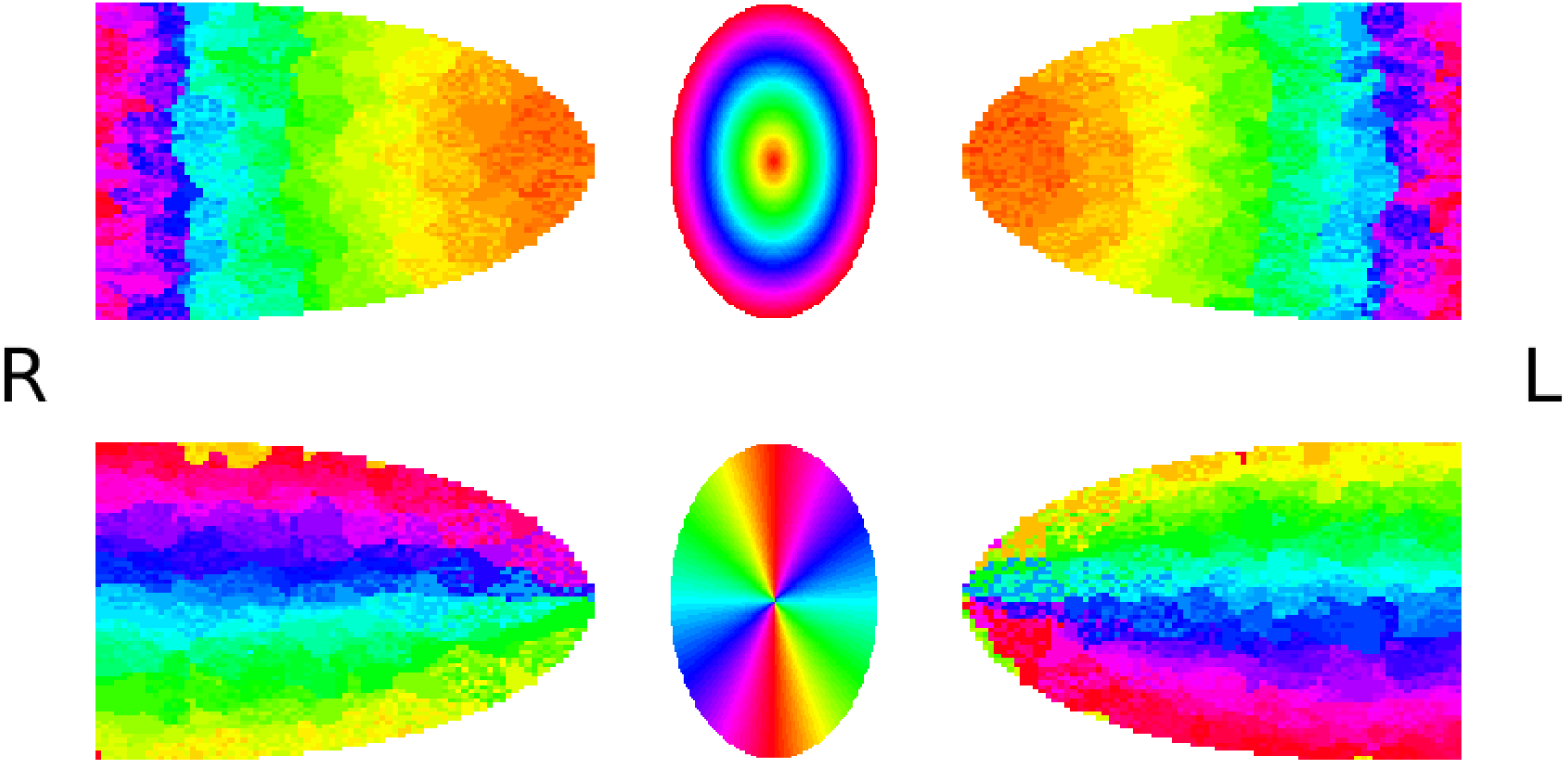
Eccentricity (top) and polar angle (bottom) maps resulting from the proposed gradient descent receptive field mapping procedure. Receptive field locations of each vertex in Cartesian cordinates are given by the location of the peak value in its weight vector and are transformed to polar coordinates.

**Figure 5.**
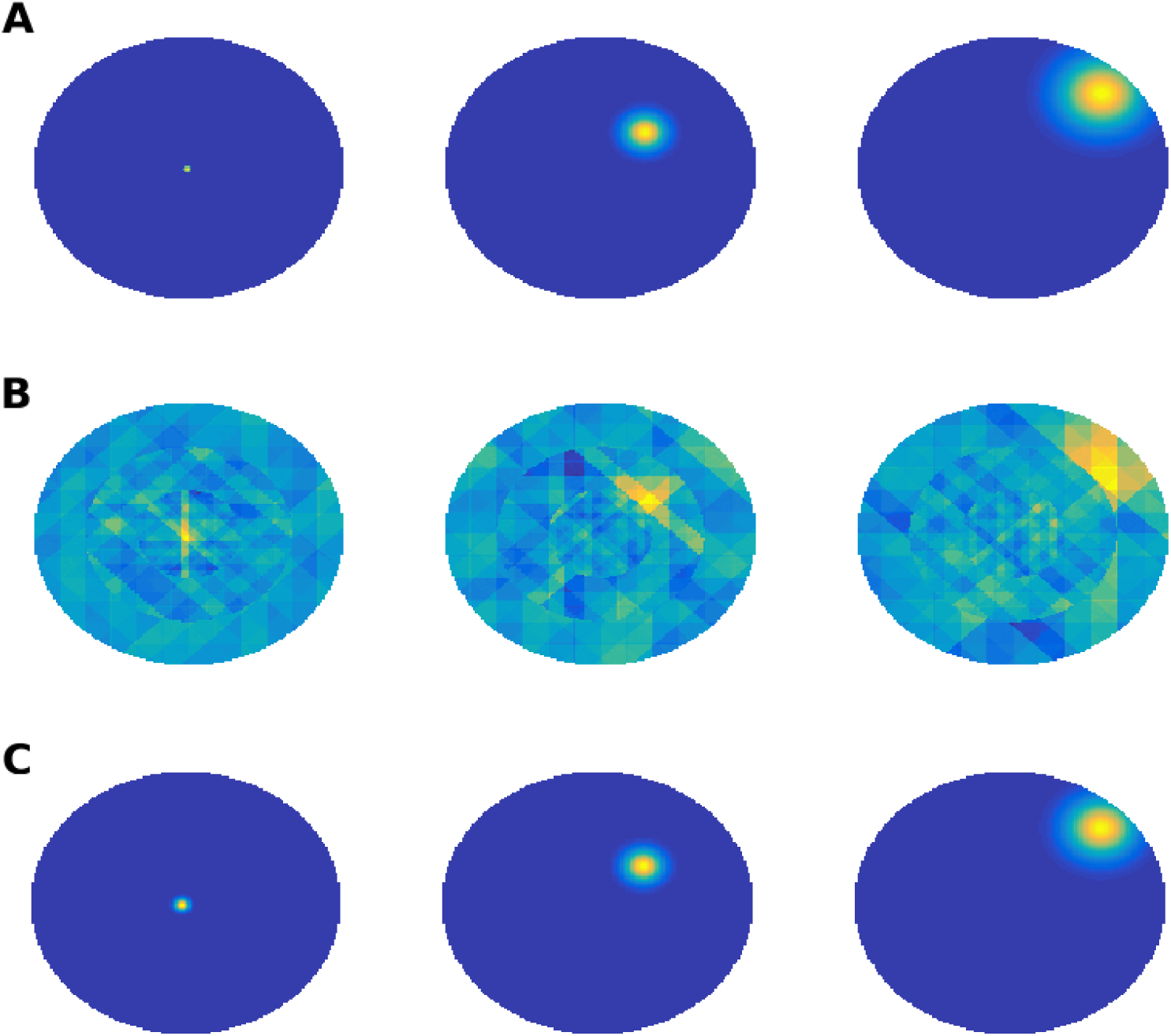
Representative receptive fields at three eccentricities (0°, 4°, and 8°). **A)** Ground-truth isotropic Gaussian receptive fields of the V1 model. **B)** Raw receptive field estimates resulting from the gradient descent mapping procedure. **C)** Receptive fields resulting from fitting an isotropic Gaussian model to the estimates.

**Figure 6.**
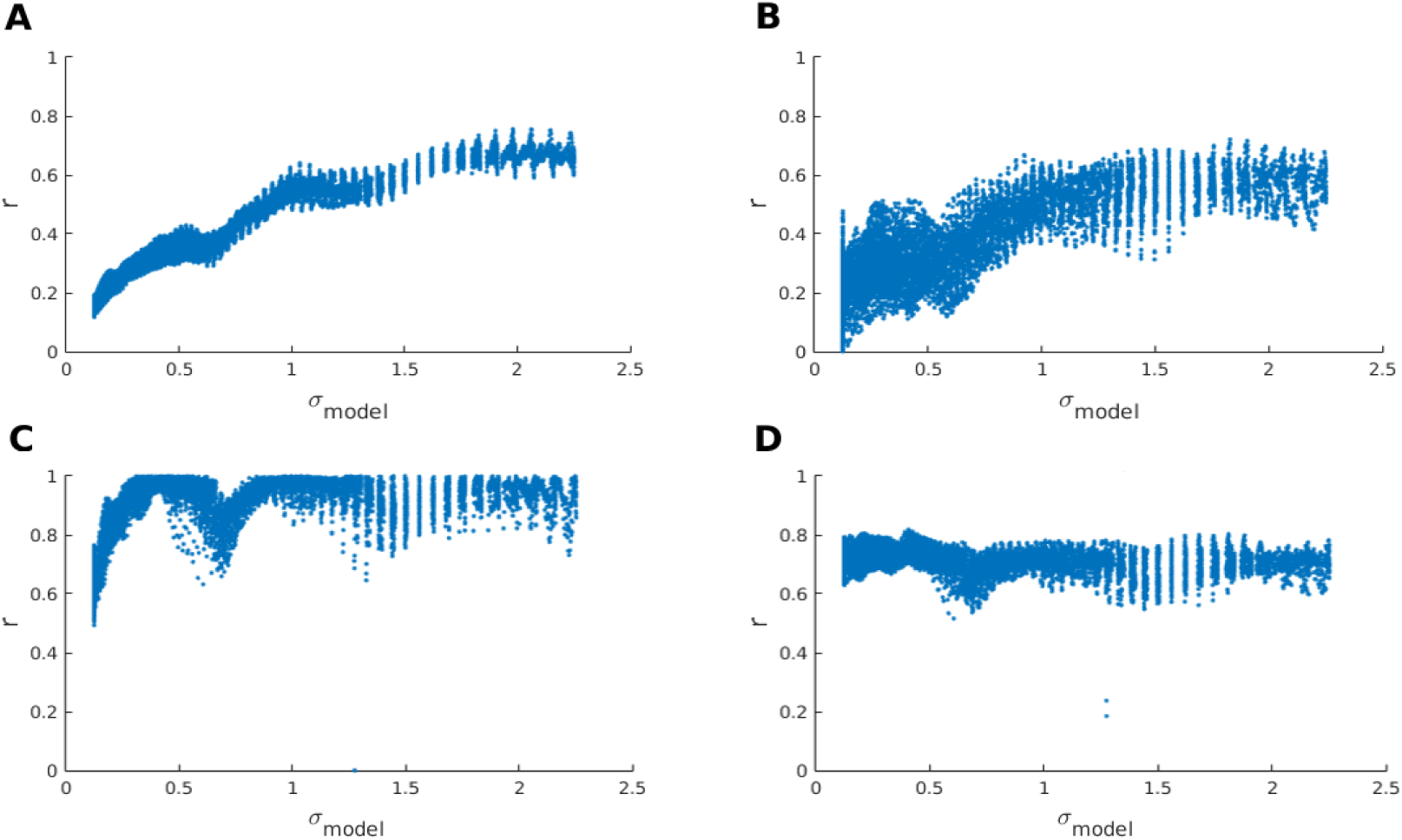
Performance as a function of ground-truth receptive field size. **A)** Correlation between model and raw estimated receptive fields for each vertex. **B)** Correlations between model BOLD activity in response to the test stimulus and predicted BOLD activity in response to the stimulus based on raw estimated receptive fields. **C)** Correlation between model and Gaussian-fit receptive field for each vertex. **D)** Correlations between model BOLD activity in response to the test stimulus and predicted BOLD activity in response to the stimulus based on Gaussian-fit receptive fields.

## Computational Performance

For the proposed mapping procedure to be practically useful in a real-time setting it is essential that all necessary computations can be carried out a rate that matches fMRI signal acquisition which typically ranges between 0.33 Hz (TR=3 s) 0.67 Hz (TR=1.5 s). The speed with which the proposed algorithm performs real-time signal analysis and updates the 22500 pixels by 11520 vertices weight matrix used here is estimated on two systems. The first is an HP(r) Z440 workstation with an Intel(r) Xeon(r) E5-1650 v3 (3.5 GHz, 15 MB cache, 6 cores) processor and 32 GB DDR4 RAM running Linux Ubuntu 14.04 as operating system. The second is a desktop computer with an Intel(r) Core i5-6600K v6 (3.5 GHz, 6 MB cache, 4 cores) processor and 16 GB DDR4 RAM running Microsoft(®) Windows 10 Professional as operating system. The algorithm has been implemented in MATLAB (2015a, The MathWorks, Natick, MA). On the first system, median and maximum code execution times are 0.51 s and 0.61 s, respectively. On the second system, median and maximum code execution times are 0.83 s and 1.08 s, respectively. Furthermore, the algorithm requires a total of 4.1 GiB of RAM, 2 GiB of which are used to store the weight matrix. These measures indicate that it is indeed feasible to use the proposed algorithm for real-time mapping of population receptive fields. Subsequent fitting of the parametric model to the weights obtained from the real-time mapping procedure takes ∼3.7 min and ∼2.9 min on the first and second system, respectively.

## Discussion

A real-time population receptive field estimation procedure based on gradient descent has been developed and evaluated in light of circularly symmetric ground-truth receptive fields in a model of V1 topography. The algorithm showed good performance in estimating receptive field location and size, robustness in the face of spatio-temporal distortions, and good computational performance. Fitting a parametric model to the raw estimates obtained from the algorithm further improves receptive field estimates and allows for accurate prediction of BOLD activity in response to new stimuli. While this fitting cannot be achieved in real-time, it is sufficiently fast to be computed during a short break for the subject (or potentially during a resting state or anatomical measurement) and to subsequently be used for any real-time applications.

It is important to note that the presented approach is merely a proof of concept and as such rather simple. However, it is vastly extendible. For instance, while in the present study the gradient descent approach is used to adjust weights linking the visual field directly to the cortex, it is possible to interpose complex filtering steps (cf. ***St-Yves and Naselaris, 2017***). That is, a fixed set of spatial filters could be continuously applied to the effective stimulus with adjustable weights linking this filter set to the cortex. In this manner, more complex response profiles of voxels could be estimated in real time.

## Acknowledgments

The author would like to thank Koen Frolichs for kindly providing the honeycomb test stimulus. The author would further like to thank Carmine Gnolo for his helpful comments.

